# Squeakr: An Exact and Approximate *k*-mer Counting System

**DOI:** 10.1101/122077

**Authors:** Prashant Pandey, Michael A. Bender, Rob Johnson, Rob Patro

## Abstract

**Motivation:** *k*-mer-based algorithms have become increasingly popular in the processing of high-throughput sequencing (HTS) data. These algorithms span the gamut of the analysis pipeline from *k*-mer counting (e.g., for estimating assembly parameters), to error correction, genome and transcriptome assembly, and even transcript quantification. Yet, these tasks often use very different *k*-mer representations and data structures. In this paper, we set forth the fundamental operations for maintaining multisets of *k*-mers and classify existing systems from a data-structural perspective. We then show how to build a *k*-mer-counting and multiset-representation system using the counting quotient filter (CQF), a feature-rich approximate membership query (AMQ) data structure. We introduce the *k*-mer-counting/querying system Squeakr (Simple Quotient filter-based Exact and Approximate Kmer Representation), which is based on the CQF. This off-the-shelf data structure turns out to be an efficient (approximate or exact) representation for sets or multisets of *k*-mers.

**Results:** Squeakr takes 2×−3;4.3× less time than the state-of-the-art to count and perform a random-point-query workload. Squeakr is memory-efficient, consuming 1.5X–4.3X less memory than the state-of-the-art. It offers competitive counting performance, and answers point queries (i.e. queries for the abundance of a particular *k*-mer) over an order-of-magnitude faster than other systems. The Squeakr representation of the *k*-mer multiset turns out to be immediately useful for downstream processing (e.g., de Bruijn graph traversal) because it supports fast queries and dynamic *k*-mer insertion, deletion, and modification.

**Availability:** https://github.com/splatlab/squeakr

**Contact**

## 1 Introduction

There has been a tremendous increase in sequencing capacity thanks to the rise of massively parallel high-throughput sequencing (HTS) technologies. Many of the new computational approaches for dealing with the increasing amounts of HTS data use the ***k-mer***—a string of *k* bases— as the atomic unit of sequence analysis. For example, most HTS-based genome and transcriptome assemblers use *k*-mers to build de Bruijn graphs (see e.g., Pevzner *et al*. (2001); Zerbino and Birney (2008); Bankevich *et al*. (2012); Simpson *et al*. (2009); Grabherr *et al*. (2011); Schulz *et al*. (2012)). De-Bruijn-graph-based assembly is favored, in part, because it eliminates the computationally burdensome “overlap” approach of the more traditional overlap-layout-consensus assembly (Koren *et al*., 2016).

*k*-mer-based methods are also heavily used for preprocessing HTS data to perform error correction (Liu *et al*., 2013; Song *et al*., 2014; Heo *et al*., 2014) and digital normalization (Brown *et al*., 2012; Zhang *et al*., 2014). Even in long-read (“3rd-generation”) sequencing-based assembly, the *k*-mer acts as a building block to help find read overlaps (Berlin *et al*., 2015; Carvalho *et al*., 2016) and to perform error correction (Salmela and Rivals, 2014; Salmela *et al*., 2016).

*k*-mer-based methods reduce the computational costs associated with many types of HTS analysis. These include transcript quantification using RNA-seq (Patro *et al*., 2014; Zhang and Wang, 2014), taxonomic classification of metagenomic reads (Wood and Salzberg, 2014; Ounit *et al*., 2015), and search-by-sequence over large repositories HTS-based sequencing experiments (Solomon and Kingsford, 2016).

Many of the analyses listed above begin by counting the number of occurrences of each *k*-mer in a sequencing dataset. In particular, *k*-mer counting is used to weed out erroneous data caused by sequencing errors. These sequencing errors most often give rise to “singleton” *k*-mers (i.e. *k*-mers that occur only once in the dataset), and the number of singletons grows linearly with the size of the underlying dataset. *k*-mer counting identifies all singletons, so that they can be removed.

*k*-mer counting is nontrivial because it needs to be done quickly, the datasets are large, and the frequency distribution of the *k*-mers is often skewed. There are many different system architectures for *k*-mer counters, because there are many different competing performance issues, including space consumption, cache-locality, and scalability with multiple threads.

However, generally, *k*-mer counters are not designed to support efficient ***point queries*,** i.e., queries for the count of an arbitrary *k*-mer. Fast point queries are helpful in downstream analysis, for example, in de Bruijn graph traversal (Chikhi and Rizk, 2013), and search (Solomon and Kingsford, 2016), and are also important for computing inner-products between datasets (Vinga and Almeida, 2003; Murray *et al*., 2016).

In general, the focus of recent *k*-mer-counting methods has been on counting performance or memory usage, with less emphasis given to query performance. Yet, many downstream analyses could benefit from a representation that supports efficient queries. Squeakr outperforms or is competitive with existing solutions in terms of counting performance and memory usage, yet provides faster queries. Most applications perform a combination of counting and querying and often querying is more prevalent than counting. Because Squeakr is so much faster for queries it results in faster overall applications with Squeakr than with other systems.

### *k*-mer Counting Systems and AMQ Data Structures

Many *k*-mer-counting approaches have been proposed in recent years, and are embodied in popular *k*-mer-counting tools such as Jellyfish (Marçais and Kingsford, 2011), BFCounter (Melsted and Pritchard, 2011), DSK (Rizk *et al*., 2013), KMC2 (Deorowicz *et al*., 2015), and Turtle (Roy *et al*., 2014).

These tools, as well as other sequence-analysis systems (Chikhi and Rizk, 2013), use the Bloom filter (Bloom, 1970) as a data-structural workhorse. The Bloom filter is a well known example of an ***approximate membership query*** (AMQ) data structure, which maintains a compact and probabilistic representation of a set or multiset. AMQs save space by allowing membership queries occasionally to return false positive answers.

*k*-mer-counting systems such as Jellyfish2 (Marçais and Kingsford, 2011), BFCounter (Melsted and Pritchard, 2011), and Turtle (Roy *et al*., 2014) use a Bloom filter to identify and filter out singleton *k*-mers, thus reducing the memory consumption. Then the systems resort to larger hash tables, or other, more traditional data structures, for the actual counting. Under such a strategy, *k*-mers are inserted into the Bloom filter upon first observation, and they are stored in a hash table (or other exact counting data structure) along with their counts upon subsequent observations.

One drawback of the Bloom filter is that it supports a relatively small set of operations and suffers from some performance issues (e.g., poor cache locality). *k*-mer-counting systems based on Bloom filters have to work around these performance and feature limitations. The limitations of the Bloom filter mean that (at least) two separate data structures need to be maintained: one for membership and one for counting. This requires all inserts to lookup and inserts in multiple structures. The Bloom filter requires a tight estimation of the number of *distinct k*-mers to get good space usage. Moreover, it does not support resizing, deletes, and counting. Additionally, a single counting quotient filter data structure is often more space efficient than a Bloom filter and hash table combination.

Furthermore, exact *k*-mer counts are often not required, and memory usage can be reduced even further, and the simplicity of the underlying algorithm improved, by replacing the Bloom filter and exact counting data structure by a single probabilistic data structure. For example, Zhang *et al*. (2014) demonstrate that the count-min sketch (Cormode and Muthukrishnan, 2005) can be used to answer *k*-mer presence and abundance queries approximately. Such approaches can yield order-of-magnitude improvements in memory usage. However, a frequency estimation data structure, like count-min sketch, can also blow up the memory usage for skewed data distributions like *k*-mers in sequencing datasets (Pandey *et al*., 2016).

There do exist more feature-rich AMQs. In particular, the counting quotient filter (CQF) (Pandey *et al*., 2016), supports operations such as insertions, deletions, counting (even on skewed datasets), resizing, merging, and highly concurrent accesses.

### Results

In this paper we show how to build a *k*-mer-counting and multiset-representation system using the recently-introduced counting quotient filter (CQF) (Pandey *et al*., 2016), a feature-rich approximate membership query (AMQ) data structure.

We show that this off-the-shelf data structure is well suited to serve as a natural and efficient structure for representing and operating on multisets of *k*-mers (exactly or approximately). We make our case by developing and evaluating a *k*-mer-counting/querying system Squeakr (Simple Quotient filter-based Exact and Approximate Kmer Representation), which is based on the CQF.

Our CQF representation is space efficient and fast, and it offers a rich set of operations. The underlying CQF is easily tuned to trade off between space and accuracy/precision, depending upon the needs of the particular application. In the application of *k*-mer counting, we observe that the counting quotient filter is particularly well suited to the highly skewed distributions that are typically observed in HTS data.

Our representation is powerful, in part, because it is dynamic. Unlike, the Bloom filter (Bloom, 1970), which is commonly used in *k*-mer-counting applications, Squeakr has the ability to modify and remove *k*-mers. Unlike the count-min sketch (Cormode and Muthukrishnan, 2005), Squeakr maintains nearly exact (or lossless) representations of the counts compactly.

In Squeakr, one can enumerate the hashes of the *k*-mers present in the structure, allowing *k*-mer multisets to be easily compared and merged. One interesting feature is that approximate multisets of different sizes can be efficiently compared and merged. This capability is likely to have other advantages beyond what we explore in this paper; for example it could be instrumental in improving the sequence Bloom tree structure used for large-scale search (Solomon and Kingsford, 2016).

We benchmark two settings of our system, Squeakr and Squeakr-exact, the latter of which supports exact counting via an invertible hash function, albeit at the cost of using more space. We compare both Squeakr and Squeakr-exact with state-of-the-art *k*-mer counting systems KMC2 and Jellyfish2. Squeakr takes 2×−4.3× less time to count and perform a random-point-query workload (de Bruijn graph graph traversal) than KMC2. Squeakr uses considerably less memory (1.5×−4.3×) than KMC2 and Jellyfish2. Squeakr offers insertion performance similar to that of KMC2 and faster than Jellyfish2. Squeakr offers an order-of-magnitude improvement in point query performance. We test the effect of query performance under both random and application-specific workloads (e.g., de Bruijn graph traversal). For point query workloads Squeakr is 3.2×−24× faster than KMC2 and Jellyfish2. For de Bruijn graph graph traversal workload Squeakr is 2× −4.3× faster than KMC2.

## 2 Methods

We begin by first describing the counting quotient filter data structure. Then, we describe the design of Squeakr; how we use the counting quotient filter in Squeakr for counting *k*-mers, and how we efficiently parallelize Squeakr to scale with multiple threads.

### 2.1 Counting quotient filter

The counting quotient filter (CQF) implements a counting filter data structure (Pandey *et al*., 2016) by storing a compact, lossless representation of the multiset *h*(*S*), where 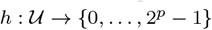 is a hash function and *S* is a multiset of items drawn from a universe 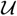. The CQF sets 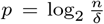 and obtains a false-positive rate *δ* while handling up to *n* insertions (Bender *et al*., 2012).

The counting quotient filter divides *h*(*x*) into its first *q* bits, ***quotient*** *h*_0_ (*x*), and its remaining *r* bits, ***remainder*** *h*_1_ (*x*). The counting quotient filter maintains an array *Q* of 2^*q*^ *r*-bit slots, each of which can hold a single remainder. When an element *x* is inserted, the counting quotient filter attempts to store the remainder *h*_1_ (*x*) in the ***home slot*** *Q*[*h*_0_ (*x*)]. If that slot is already in use, then the counting quotient filter uses a variant of linear probing (using few metadata-bits per slot), to find an unused slot where it can store *h*_1_ (*x*) (Pandey *et al*., 2016).

Instead of storing multiple copies of the same item to count, like a quotient filter, the counting quotient filter employs an encoding scheme to count the multiplicity of items. The encoding scheme enables the counting quotient filter to maintain *variable-sized* counters. This is achieved by using slots originally reserved to store the remainders to, instead, store count information. The metadata bits maintained by the counting quotient filter allows this dynamic reuse of remainder slots for large counters while still ensuring the correctness of all counting quotient filter operations.

The variable-sized counters in the counting quotient filter enable the data structure to handle highly skewed datasets efficiently. By reusing the allocated space, the counting quotient filter avoids wasting extra space on counters and naturally and dynamically adapts to the frequency distribution of the input data. The counting quotient filter never takes more space than a quotient filter for storing the same multiset. For highly skewed distributions, like those observed in HTS-based datasets, it occupies only a small fraction of the space that would be required by a comparable (in terms of false-positive rate) quotient filter.

### 2.2 Squeakr design

A *k*-mer counting system starts by reading and parsing the input file(s) (i.e., `FASTA/FASTQ` files) and extracting reads. These reads are then processed from left to right, extracting each read’s constituent *k*-mers. The *k*-mers may be considered as they appear in the read, or, they may first be *canonicalized* (i.e., a *k*-mer is converted to the lexicographically smaller of the original *k*-mer and its reverse-complement). The goal of a *k*-mer counting system is to count the number of occurrences of each *k*-mer (or canonical *k*-mer) present in the input dataset.

Squeakr has a relatively simple design compared to many existing *k*-mer counting systems. Many *k*-mer counting systems use multiple data structures, e.g., an Approximate Membership Query (AMQ) data structure to maintain all the singletons and a hash table (Marçais and Kingsford, 2011) or compaction-and-sort based data structure (Roy *et al*., 2014) for the actual counting. The motivation behind using multiple data structures is primarily to reduce the memory requirements. Yet, having to maintain and modify multiple data structures when processing each *k*-mer can slow the counting process, and add complexity to the design of the *k*-mer counter. Other *k*-mer counting systems use domain specific optimizations (e.g., minimizers (Roberts *et al*., 2004)) to achieve faster performance (Roberts *et al*., 2004; Deorowicz *et al*., 2015).

Squeakr uses a single, off-the-shelf data structure, the counting quotient filter, with a straightforward system design, yet still offers superior performance in terms of memory and running and query time.

In Squeakr we have a single-phase process for counting *k*-mers in a read data set. Each thread performs the same set of operations; reading data from disk, parsing and extracting *k*-mers, and inserting *k*-mers in the counting quotient filter. The input file is read in chunks of 16MB, and each chunk is then parsed to find the last complete read record.

To synchronize operations among multiple threads, Squeakr uses a lock-free queue (Boost, 2014) for storing the state of each file being read from disk, and a *thread-safe* counting quotient filter for inserting *k*-mers. Each thread executes a loop in which it grabs a file off the lock-free queue, reads the next 16MB chunk from the file^1^, returns the file to the lock-free queue, and then parses the reads in the 16MB chunk and inserts the resulting *k*-mers into a shared, thread-safe quotient filter. This approach enables Squeakr to parallelize file reading and parsing, improving its ability to scale with more threads. KMC2, on the other hand, uses a single thread for each file to read and decompress which can sometimes become a bottleneck e.g., when one input file is much larger than others.

Squeakr uses a thread-safe counting quotient filter (Pandey *et al*., 2016) for synchronizing insert operations among multiple threads. Using a thread-safe counting quotient filter, multiple threads can simultaneously insert *k*-mers into the data structure. Each thread acquires a lock on the region where the *k*-mer must be inserted and releases the lock once it is done inserting the *k*-mer. *k*-mers that hash to different regions of the counting quotient filter may be inserted concurrently.

This scheme of using a single, thread-safe counting quotient filter scales well with an increasing number of threads for smaller datasets, e.g., *F. vesca* and *G. gallus* (see Table 1) where the skewness (in terms of *k*-mer multiplicity) is not very high. However, for larger highly-skewed datasets, the thread-safe counting quotient filter scheme does not scale well. This is due, in part, to the fact that these data have many highly-repetitive *k*-mers, causing multiple threads to attempt to acquire the same locks. This results in excessive lock contention among threads trying to insert *k*-mers, and prevents an increase in the number of threads from leading to a concordant decrease in the total time required to count all *k*-mers.

**Table 1.** datasets used in the experiments. The file size is in GB. All the datasets are compressed with gzip compression.

| dataset | File size | #Files | #$k$-mer instances | #Distinct $k$-mers |
| --- | --- | --- | --- | --- |
| *F. vesca* | 3.3 | 11 | 4134078256 | 632436468 |
| *G. gallus* | 25.0 | 15 | 25337974831 | 2727529829 |
| *M. balbisiana* | 46.0 | 2 | 41063145194 | 965691662 |
| *H. sapiens 1* | 67.0 | 6 | 62837392588 | 6353512803 |
| *H. sapiens 2* | 99.0 | 48 | 98892620173 | 6634382141 |

**Scaling with multiple threads.** Large datasets with high skewness contain *k*-mers with very high multiplicity (of the order of billions). Such *k*-mers causes hotspots, and lead to excessive lock contention among threads in the counting quotient filter. To overcome the issue of excessive lock contention, Squeakr tries to reduce the time spent by threads waiting on a lock by amortizing the cost of acquiring a lock.

As explained in (Pandey *et al*., 2016), the time spent by threads while waiting for a lock can be utilized to do local work. As shown in Figure 1, we assign a local counting quotient filter to each thread. Now, during insertion, each thread first tries to insert the *k*-mer in the thread-safe, global counting quotient filter. If the thread acquires the lock in the first attempt, then it inserts the *k*-mer and releases the lock. Otherwise, instead of waiting on the lock to be released, it inserts the *k*-mer in the local counting quotient filter and continues. Once the local counting quotient filter becomes full, the thread dumps the *k*-mers present in the local counting quotient filter into the global counting quotient filter before processing any new *k*-mers.

**Fig. 1:**
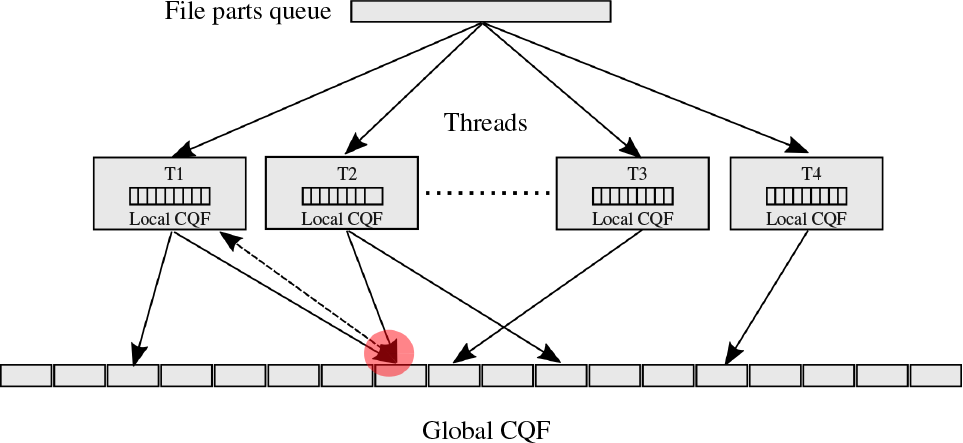
Squeakr system design: each thread has a local CQF and there is a global CQF. The dotted arrow shows that one thread did not get the lock in the first attempt and had to insert the item in the local CQF.

The above approach helps to reduce the time spent by threads while waiting on a lock and also amortizes the cost of acquiring a lock. Intuitively, repetitive *k*-mers are the ones for which it is hardest to acquire the lock in the global counting quotient filter. When a thread encounters such a *k*-mer and fails to obtain the corresponding lock in the global counting quotient filter, the thread instead immediately inserts those *k*-mers in the local (lockless) counting quotient filter and continues processing data. Moreover, these repetitive *k*-mers are first counted in the local counting quotient filter before being inserted with their corresponding counts in the global counting quotient filter. Under this design, instead of inserting multiple instances of the *k*-mer in the global counting quotient filter, requiring multiple acquisitions of a global counting quotient filter lock, Squeakr only insert the *k*-mers a few times with their counts aggregated via the local counting quotient filters.

In Squeakr even while maintaining multiple data structures, a local counting quotient filter per thread and a global counting quotient filter, one operation is performed for the vast majority of *k*-mers. While inserting *k*-mers that occur only a small number of times, threads obtain the corresponding lock in the global counting quotient filter in the first attempt. These *k*-mers are only inserted once. On the other hand, for repetitive *k*-mers, instead of acquiring the lock for each observation, we insert them in the local counting quotient filter and only insert them into the global counting quotient filter once the local counting quotient filter gets full.

### 2.3 Squeakr is both an approximate and exact *k*-mer counter

Squeakr is capable of acting as either an approximate or an exact *k*-mer counter. In fact, this can be achieved with no fundamental changes to the underlying system; but simply by increasing the space dedicated to storing each hash’s remainder, and by adopting an invertible hash function.

In Squeakr each *k*-mer in the read dataset is represented as a bit vector using 2*k* bits, i.e., each base-pair is represented using 2 bits. As explained in Section 2.1, to achieve a maximum allowable false-positive rate δ the counting quotient filter requires a *p*-bit hash function, where 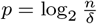 and *n* is the number of distinct *k*-mers. For example, to achieve a false-positive rate of 1/512 for a dataset with 2^30^ distinct *k*-mers, we need a 39-bit hash function. In Squeakr, we use the Murmur hash function (Appleby, 2016), by default, for hashing *k*-mers.

In the counting quotient filter, the *p*-bit hash is divided into *q* quotient bits and r remainder bits. The maximum false-positive rate is bounded by 2^−*r*^ (Bender *et al*., 2012). In Squeakr, we assign *q* = log *n*, where *n* is the number of distinct *k*-mers in the dataset and we use 9-bit remainders to achieve a false-positive rate of 1/512.

In order to convert Squeakr from an approximate *k*-mer counter to an exact *k*-mer counter, we need to use a *p*-bit *invertible* hash function, where *p* = 2*k*. In Squeakr-exact, we use the Inthash hash function (Li, 2016) for hashing *k*-mers. For a dataset with *n* distinct *k*-mers and a *p*-bit hash function, the remainder *r* = *p* – log_2_ *n*. For example, for *n* = 2^30^ and *k* = 28 (i.e., *p* = 56), we need *r* = 26 bits. This is still far less than the 56 bits that would be required to store each *k*-mer key explicitly.

## 3 Results

In this section we evaluate our implementations of Squeakr and Squeakr-exact. Squeakr-exact, as described in Section 2.3, is an exact *k*-mer counting system that uses the counting quotient filter with a *p*-bit invertible hash function, where *p* is the number of bits to represent a *k*-mer in binary. Squeakr is an approximate *k*-mer counting system that also uses the counting quotient filter but takes much less space than Squeakr-exact. The space savings comes from the fact that Squeakr allows a very small false-positive rate.

We compare both versions of Squeakr with state-of-the-art *k*-mer counting systems in terms of speed, memory efficiency, and scalability with multiple threads. We compare Squeakr against two *k*-mer counting systems; KMC2 (Danek, 2016) and Jellyfish2 (Marçais and Kingsford, 2011). KMC2 is currently the fastest *k*-mer counting system (Deorowicz *et al*., 2015), although not the most frugal in terms of memory usage (when not run in disk-based mode). Jellyfish2, though not the fastest or most memory-frugal system, is very widely used, and internally uses a domain specific hash-table to count *k*-mers, and is thus methodologically similar to Squeakr.

Khmer (Zhang *et al*., 2014) is the only approximate multiset representation and uses a count-min sketch. Here, we don’t compare Squeakr against Khmer, since they are geared toward somewhat different use-cases. Squeakr exhibits a very small error rate, and is intended to be used in places where one might otherwise use an exact *k*-mer counter, while Khmer is designed much more as a sketch, to perform operations on streams of *k*-mers for which near-exact counts are not required.

Squeakr is an in-memory *k*-mer counter and we compare it against other in-memory *k*-mer counting systems. We currently only support *k*-mers of maximum length 32, though, it is not a fundamental limitation of the counting quotient filter.

We evaluate each system on two fundamental operations, counting and querying. We use multiple datasets to evaluate counting performance, and a subset of those datasets for query performance. We evaluate queries for existing *k*-mers and absent *k*-mers (uniformly random *k*-mers) in the dataset. We also evaluate Squeakr for performing queries for *k*-mers as they appear in the context of de Bruijn graph traversal. Traversing the de Bruijn graph is a critical step in any De-Bruijn-graph-based assembly, and using an AMQ for a compact representation and fast traversal of the de Bruijn graph has been shown in the past (Pell *et al*., 2012). To evaluate the ability of the counting quotient filter to represent a de Bruijn graph and deliver fast query performance during traversal, we performed a benchmark where we query *k*-mers as they appear in the de Bruijn graph.

Other than counting and queries, we also evaluate Squeakr for computing the inner-product between the *k*-mer abundance vectors of a pair of datasets. The comparison of the *k*-mer composition of two different strings (or entire datasets) has proven a fruitful approach for quickly and robustly estimating their overall similarity. In fact, many methods exist for the so-called alignment-free comparison of sequencing data (Vinga and Almeida, 2003). Recently, (Murray *et al*., 2016) introduced the *k*-mer weighted inner product as an estimate of the similarity between genomic/metagenomic datasets. Prior work suggests that ability to compute fast inner-product between two datasets is important. To assess the utility of the CQF in enabling such types of comparisons, we performed a benchmark to evaluate the performance of Squeakr to compute the inner-product (or cosine-similarity) between two datasets.

### 3.1 Experimental Setup

Each system is tested for 28-mers and all the experiments are performed in-memory. We use several datasets for our experiments, which are listed in Table 1. All the experiments were performed on an Intel(R) Xeon(R) CPU (E5−2699 v4 @ 2.20GHz with 44 cores and 56MB L3 cache) with 512GB RAM and a 4TB TOSHIBA MG03ACA4 ATA HDD. In order to evaluate the scalability of the systems with multiple threads, we have reported numbers with 8 and 16 threads for all the systems and for each dataset. In all our experiments, the counting quotient filter was configured with a maximum allowable false-positive rate of 1 /512. For reporting time and memory metrics, we have taken the average over two runs for all the benchmarks. The time reported in all the benchmarks is in seconds and memory (RAM) is in GBs.

**Counting benchmarks.** For each dataset we have only counted the canonical *k*-mers. In order to isolate the counting performance of each system, we have only reported the time taken by the system to count *k*-mers excluding the initialization time (e.g., allocating data structures, locks, etc), though a small fraction of the total time. The time reported for counting benchmarks is the total time taken by the system to read data off disk, parse it, count *k*-mers, and write the *k*-mer representation to disk.

The memory reported is the maximum RAM required by the system while counting *k*-mers, as given by “/usr/bin/time”. The RAM mentioned in Table 2 is the average RAM required by the system for 8 and 16 threads. Both Squeakr and Jellyfish2 require to give as a parameter the number of distinct *k*-mers in the dataset. Squeakr needs the number of distinct *k*-mers (approximate to next closet power of 2) as an input. Squeakr takes the approximation of number of distinct *k*-mers as the number of slots to create the CQF. We used Mohamadi *et al*. (2017) to estimate the number of distinct *k*-mers in datasets. As explained in Section 2.2, KMC2 can be bottlenecked to decompress bzip2 compressed files. Therefore, we compress all the files to gzip. In our experiments gzip decompression was fast enough and never looked as a bottleneck.

**Table 2.** Amount of RAM used by KMC2, Squeakr, and Jellyfish2 for various datasets for in-memory experiments. RAM is in GB.

| dataset | KMC2 | Squeakr | Squeakr-exact | Jellyfish2 |
| --- | --- | --- | --- | --- |
| *F. vesca* | 8.3 | 4.8 | 9.3 | 8.3 |
| *G. gallus* | 32.8 | 13.0 | 28.8 | 31.7 |
| *M. balbisiana* | 48.3 | 11.1 | 14.2 | 16.3 |
| *H. sapiens 1* | 71.4 | 22.1 | 51.5 | 61.8 |
| *H. sapiens 2* | 107.4 | 30.8 | 60.1 | 61.8 |

**Query benchmarks.** We performed three different benchmarks for queries. First, we randomly queried for *k*-mers that we knew existed in the dataset. Second, we queried for 1.5B uniformly random *k*-mers (i.e., uniformly random 28-mers), most of which are highly unlikely to exist in the dataset. Third, we performed a de Bruijn graph traversal, walking the paths in the de Bruijn graph and querying the neighbors of each node. We have performed the query benchmarks on two different datasets, *G. gallus* and *M. balbisiana*. Also, we excluded Jellyfish2 in the de Bruijn graph benchmark because Jellyfish2’s random query performance was very slow for the first two query benchmarks. We note here that this appears to be a result of the fact that Jellyfish2 uses a sorted, compacted list to represent the final counts for each *k*-mer, rather than the hash table that is used during counting. This helps to minimize on-disk space, but results in logarithmic random query times.

In the de Bruijn graph traversal benchmark, for each distinct *k*-mer in the dataset, we traverse in the graph to find the longest, non-branching path (the longest path that can be traversed before hitting a fork). That is, for each *k*-mer, we take the suffix of length *k* **–** 1 and append each of the four possible bases to generate four new *k*-mers. Then we perform four separate queries for these newly generated *k*-mers in the database. If there is more than one newly generated *k*-mer present (i.e., a fork) then we stop. Otherwise, we continue this process. At the end, we report the total time taken and the longest path in the de Bruijn graph.

In the inner-product query benchmark, we first count the *k*-mers from two datasets and store the *k*-mer representations on disk. We then compute the inner-product between the two datasets by querying *k*-mers from one dataset in the other dataset. For this benchmark we do not load the whole representation in memory. Instead we *mmap* the representation and allow the kernel to load the respective blocks as we proceed in computing the inner-product. We excluded KMC2 and Jellyfish2 from the inner-product query benchmark because both KMC2 and Jellyfish2 do not expose an API for computing inner-product.

For all query benchmarks, we only report the time to query *k*-mers in the database. We exclude the time to read *k*-mers from an input file or to generate *k*-mers (for uniformly random query benchmark). Also, we first load the database completely in memory for all the systems before performing any queries. In case of KMC2 we load the database in random-access mode. For the de Bruijn graph traversal queries, we generate *k*-mers to traverse the graph on-the-fly. The time reported for the de Bruijn graph traversal query includes the time to generate these *k*-mers.

### 3.2 Memory requirement

Table 2 shows the maximum memory required by KMC2, Squeakr, Squeakr-exact, and Jellyfish2 for counting *k*-mers from different datasets. Squeakr requires the least RAM compared to the other systems. Even for the human datasets, Squeakr is very frugal in memory usage and completes the experiment in ≈ 30GB of RAM. Across all datasets, Squeakr takes 1.5X—4.3X less RAM than KMC2 (in in-memory mode).

Squeakr-exact takes less RAM than KMC2 for all (except *F. vesca)* datasets and Jellyfish2 for human datasets. For smaller datasets, Squeakr-exact takes approximately the same amount of RAM as Jellyfish2.

### 3.3 Counting performance

Figure 2 shows the time taken by different systems to count the *k*-mers present in the datasets.

**Fig. 2:**
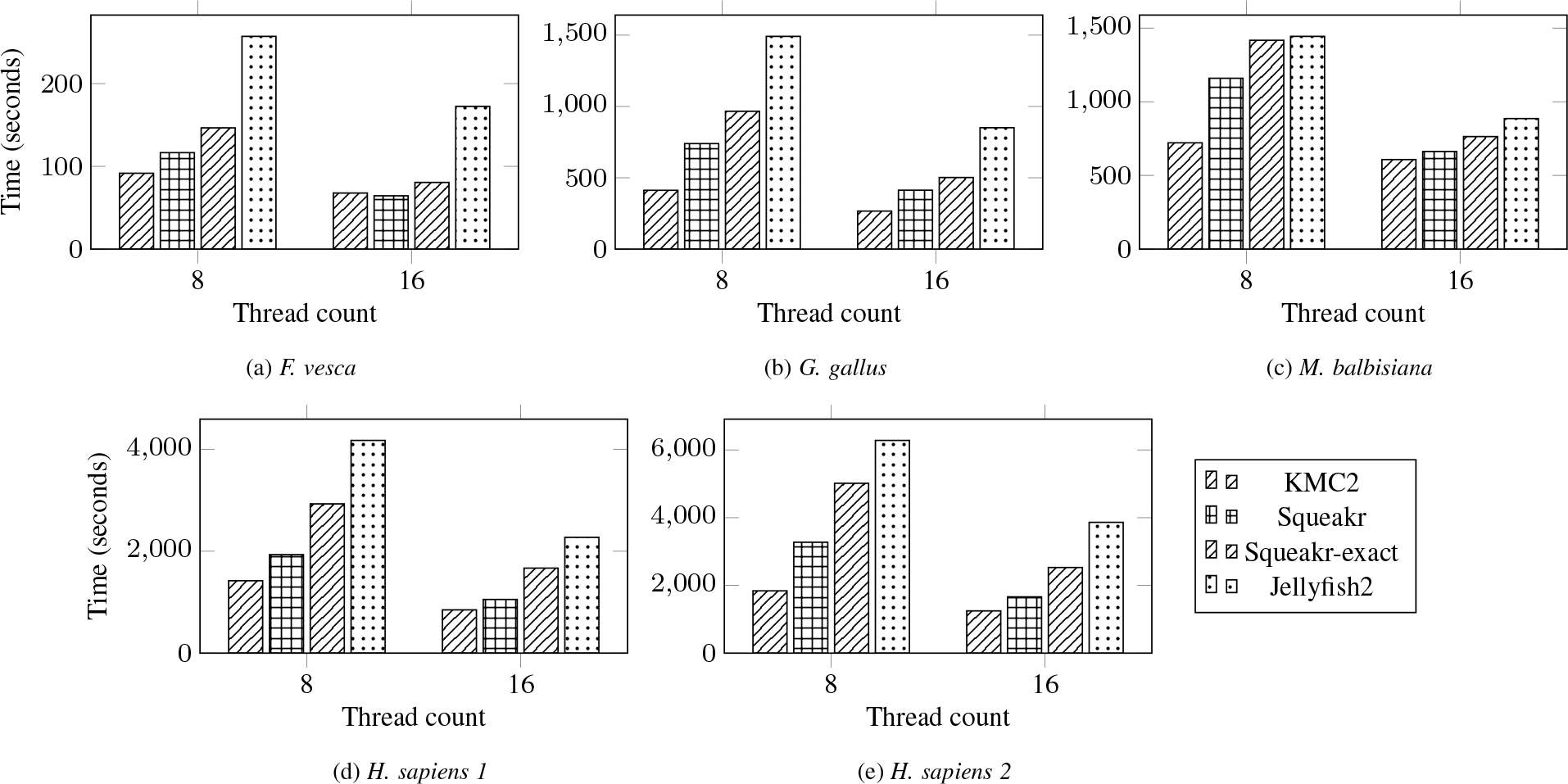
*k*-mer counting performance of KMC2, Squeakr, Squeakr-exact, and Jellyfish2 on different datasets. (Lower is better.)

KMC2 is the fastest *k*-mer counter across all datasets. Squeakr is the second fastest *k*-mer counter. For human datasets, Squeakr is 24%—43% slower than KMC2. For other datasets, Squeakr is 8%—56% slower than KMC2 and 5% faster than KMC2 for the *f.vesca* dataset.

Squeakr scales better than KMC2 with an increasing number of threads. For the *M. balbisiana* dataset, when going from 4 to 16 threads, KMC2’s time decreases by 15%, whereas Squeakr’s time decreases by 75%. For *G. gallus* dataset, when going from 4 to 16 threads, KMC2’s time decreases by 54% whereas Squeakr’s time deceases by 79%.

Squeakr-exact is slower than Squeakr for all datasets tested. However, we find that it is always faster than Jellyfish2.

### 3.4 Query performance

**Random query for existing** *k***-mers.** Figure 3a shows the random query performance for existing *k*-mers. Squeakr is 3.2X—4.9X faster than KMC2 for random queries for existing *k*-mers. Jellyfish2 is the slowest. This is likely because the on-disk representation used by Jellyfish2 is a compacted, sorted list, not the hash table used during the counting phase.

**Fig. 3:**
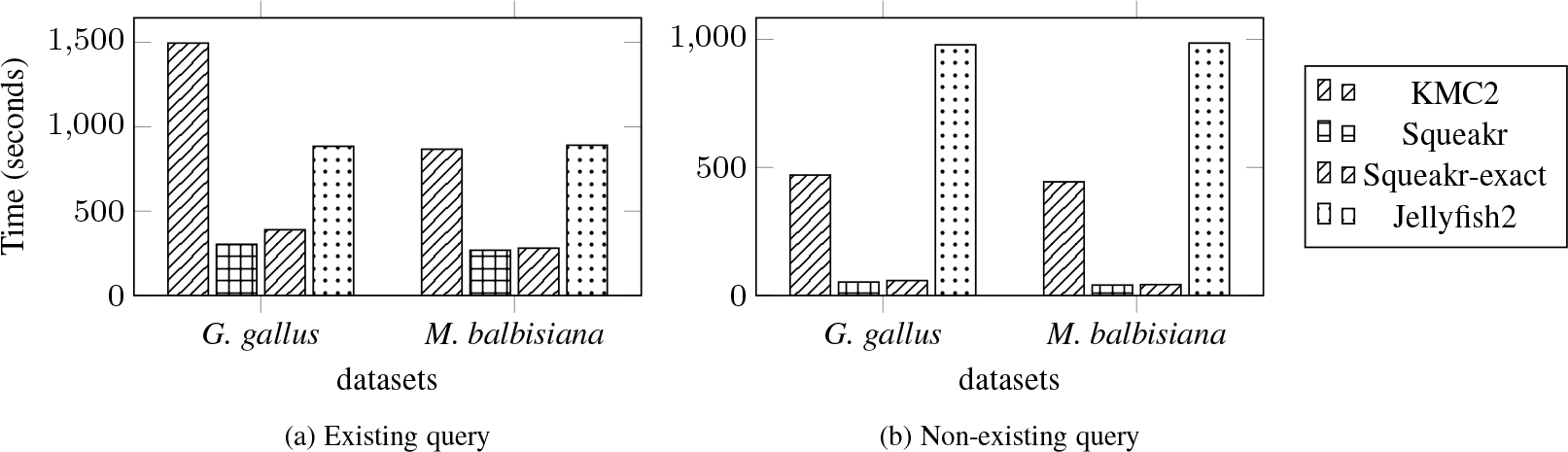
Random query performance of KMC2, Squeakr, Squeakr-exact, and Jellyfish2 on two different datasets. (Lower is better.)

**Random query for uniformly-random *k*-mers**. Figure 3b shows the random query performance for uniformly-random *k*-mers. For uniformly-random *k*-mers, Squeakr is 8.9X—10.8X faster than KMC2. Squeakr is even faster for uniformly-random queries than when querying for existing *k*-mers because there is a fast path for non-existing items in the counting quotient filter. For non-existing items, the counting quotient filter often returns the result by examining a single bit. Jellyfish2 is the slowest among the three for uniformly-random *k*-mer queries.

Both Squeakr and Squeakr-exact have similar query performance, with Squeakr-exact being slightly slower because the exact version requires a larger counting quotient filter structure.

We also evaluated the empirical false-positive rate of Squeakr, which we find to be very close to the theoretical false-positive rate. As mentioned in Section 3.1, the theoretical false-positive rate is 1/512 i.e., 0.001953125. The empirical false-positive rate reported during the benchmark is 0.0012414.

**de Bruijn graph traversal.** Table 3 shows the de Bruijn graph traversal performance of Squeakr and KMC2.

**Table 3.**
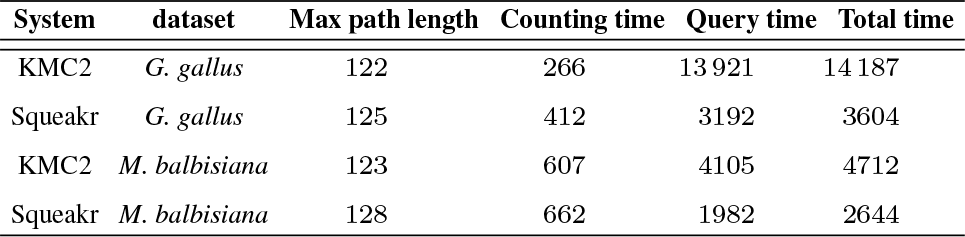
de Bruijn graph query performance on different datasets. The counting time is calculated using 16 threads. The query time is calculated using a single thread. Time is in seconds. We excluded Jellyfish2 from this benchmark because Jellyfish2 performs slowly compared to KMC2 and Squeakr for both counting and query (random query and existing *k*-mer query).

Squeakr being much faster than KMC2 for both type of queries, existing *k*-mers and non-existing *k*-mers, performs 2X—4.3X faster than KMC2 for de Bruijn graph traversal queries. For the de Bruijn graph traversal, we perform 4 queries for each *k*-mer. To continue on the path, only one out of the four queries should return true. In the whole benchmark, ≈ 75% of the queries are false queries.

Table 3 also reports the longest path present in the graph reported by both the systems. Squeakr, being an approximate *k*-mer counter has some false-positives. The length of the longest path reported by Squeakr is off from the exact answer by only 3 and 5 bases for the *G. gallus* and *M. balbisiana* data sets, respectively.

In the table, we also present the time taken to count *k*-mers in the dataset and the total time (i.e., counting time and de Bruijn graph traversal time). Squeakr is 1.7X—3.9X faster than KMC2 in terms of total time.

### 3.5 Inner-product queries

For inner-product queries, we first counted *k*-mers from two different datasets, each having ≈ 20 Billion *k*-mer instances and ≈ 965 Million distinct *k*-mers and stored the databases on disk. We then computed the inner-product between the two datasets by reading the database from disk. It took ≈ 46 seconds for Squeakr to compute the inner-product between these two datasets. This suggests that the CQF *k*-mer multiset representation provides a fast and efficient way to enumerate and query *k*-mers. This feature can be used for large-scale comparison and organization of sequencing datasets.

KMC2 has an API to perform intersection on two *k*-mer representations. We also implemented an API in the counting quotient filter for computing intersection on two on-disk counting quotient filters similar to inner-product. Both KMC2 and Squeakr take similar time (≈ 200) seconds to perform intersection on two different datasets used for the inner-product query benchmark.

## 4 Conclusion

We argue that the counting quotient filter can serve as a memory-efficient, fast, and feature-rich representation of *k*-mer multisets. We demonstrate these qualities by building a counting quotient filter-based *k*-mercountingsystem, Squeakr. Despite its relatively straightforward use of an off-the-shelf data structure, Squeakr offers great counting performance and exceptional query performance.

Squeakr is much more space efficient than other *k*-mer-counting solutions (even when representing the *k*-mer multiset exactly), and scales with multiple threads. Squeakr’s *k*-mer representation can be further used to store and traverse the weighted de Bruijn graph over the *k*-mers with high query throughput, an order-of-magnitude faster than other systems like KMC2 and Jellyfish2. Furthermore, Squeakr is dynamic, since *k*-mers can be added or removed, and their counts can be updated.

We believe that these properties make Squeakr’s *k*-mer representation a strong candidate for many downstream analysis tasks. For example, the fast query performance and ability to accurately record *k*-mer counts makes the counting quotient filter an enticing candidate atop which to build a de Bruijn graph-based transcriptome assembler. We anticipate that, with some domain specific optimizations, Squeakr’s counting quotient filter can be made even more memory efficient and can be used to efficiently store an exact weighted de Bruijn graph.

## 5 Acknowledgments

We gratefully acknowledge support from NSF grants BBSRC-NSF/BIO-1564917, IIS-1247726, IIS-1251137, CNS-1408695, CCF-1439084, and CCF-1617618, and from Sandia National Laboratories.

## Footnotes

1 Threads actually read slightly less than 16MB, since threads always break chunks at inter-read boundaries.

